# Alterations of human lung and gut microbiome in non-small cell lung carcinomas and distant metastasis

**DOI:** 10.1101/2020.01.06.895490

**Authors:** Hui Lu, Na L. Gao, Chunhua Wei, Jiaojiao Wang, Fan Tong, Huanhuan Li, Ruiguang Zhang, Hong Ma, Nong Yang, Yongchang Zhang, Ye Wang, Zhiwen Liang, Hao Zeng, Wei-Hua Chen, Xiaorong Dong

**Author notes:** Correspondence should be addressed to Wei-Hua Chen or Xiaorong Dong. Contributed equally to this work (Hui Lu, Na L. Gao, Chunhua Wei).

## Abstract

**Background:** Non-small cell lung cancer (NSCLC) is the leading cause of cancer-related deaths worldwide. Although dysbiosis of lung and gut microbiota have been associated with NSCLC, their relative contributions are unclear; in addition, their roles in distant metastasis (DM) are still illusive.

**Results:** We surveyed the fecal and sputum (as a proxy for lung) microbiota in healthy controls and NSCLC patients of various stages, and found significant perturbations of gut- and sputum-microbiota in patients with NSCLC and DM. Machine-learning models combining both microbiota (mixed models) performed better than either dataset in patient stratification, with the highest area under the curve (AUC) value of 0.842. Sputum-microbiota contributed more than the gut in the mixed models; in addition, sputum-only models performed similarly to the mixed models in most cases. Several microbial-biomarkers were shared by both microbiota, indicating their similar roles at distinct body sites. Microbial-biomarkers of distinct disease stages were mostly shared, suggesting biomarkers for distant metastasis could be acquired early. Furthermore, *Pseudomonas aeruginosa*, a species previously associated with wound infections, was significantly more abundant in brain metastasis, indicating distinct types of DMs could have different microbial-biomarkers.

**Conclusion:** Our results indicate that alterations of sputum-microbiota have stronger relationships with NSCLC and distant metastasis than the gut, and strongly support the feasibility of metagenome-based non-invasive disease diagnosis and risk evaluation.

## Background

Lung cancer (LC) is the leading cause of cancer-related deaths mortality worldwide, with non-small cell lung cancer (NSCLC) being the most common form of LC [1]. Despite the recent development of therapies for NSCLC, tumor metastasis is the main cause of recurrence and mortality in patients with NSCLC [1]. One of the key challenges is the low heritability of lung cancer susceptibility revealed by genetic studies: although numerous studies have established the important roles of somatic mutations as well as inheritable familial risks [2, 3], the genetic influence can only explain 3∼15% of the heritability [4, 5], depending on the surveyed population.

Conversely, non-genetic factors, including life styles, environmental factors and lung and gut microbes are believed to contribute mostly to the disease. Especially, numerous recent studies have shown that both lung and gut microbiota are involved in the development of LC [6–8]. For example, researchers have used samples from bronchoalveolar fluid (BALF), tissues and spontaneous sputum of lung cancer patients for bacterial identification and microbiome characterization [7, 9–11]. When compared with healthy controls, researchers have identified certain lung or oral taxa, including *Streptococcus* and *Veillonella* were enriched in the patients, which might promote LC development through inflammation and/or unappreciated mechanisms [7, 12].

In addition, dysbiosis of gut microbiome has also been associated with many cancers [8, 13, 14], including LC [8]. A previous study suggested an increase in *Enterococcus* in the stool of patients with LC, compared with the stool of healthy subjects, and a decrease in *Bifidobacterium* and *Actinobacteria* [6], which others have shown that the response to immunotherapy (IO) in NSCLC patients is associated with changes of individual species such as *Alistipes putredinis, Bifidobacterium longum* and *Prevotella copri* as well as the overall diversity of the gut microbiome [7, 8]. Furthermore, increasing evidence have shown that the gut microbiome may play important roles in cancer by modulating inflammation [15], host immune response [16, 17] and directly interacting with therapeutic drugs [18].

Despite these significant advances, two important questions remain. First, it is still unclear which microbiota has stronger association with the development of NSCLC; the relative importance of local (i.e. lung-associated) versus gut microbiota has been recently discussed [19], but no direct evidence has been provided so far. Second, their alterations along with distant metastasis of NSCLC are yet to be characterized. To address these issues, we first conducted a comprehensive survey on both fecal and sputum (as a proxy for lung) microbiota in NSCLC patients of various stages, including stage Ⅳ patients suffered from distant metastasis (DM), and compared them with healthy controls of matching demographic and clinical characteristics. We then built mathematical models using the taxonomic profiles of both gut and sputum microbiota to test their ability to distinguish patients of different disease stages and from healthy controls, and evaluate their relative contributions to the models.

## Results

### Differential microbial diversity between sputum and gut microbiotas

We enrolled in total 121 individuals who completed our study protocol (see Methods). Among which, 87 were newly diagnosed with NSCLC who had not previously received any anticancer therapy nor treated with any antibiotics, while 34 were healthy volunteers. We classified patients into distinct disease stages (i.e. from I to IV) according to the 8th American Joint Committee on Cancer (AJCC) guidelines [20]. All subjects currently lived in Hubei Province, China. As shown in Table 1, we found comparable demographic and clinical characteristics of these subjects between groups we were interested in. In this study, we used “Control”, “NSCLC”, “I_III” and “DM” to refer healthy controls, patients of all stages, patients of stages I to III and patients with distal metastasis (DM, also referred as to stage IV), respectively.

**Table 1.**
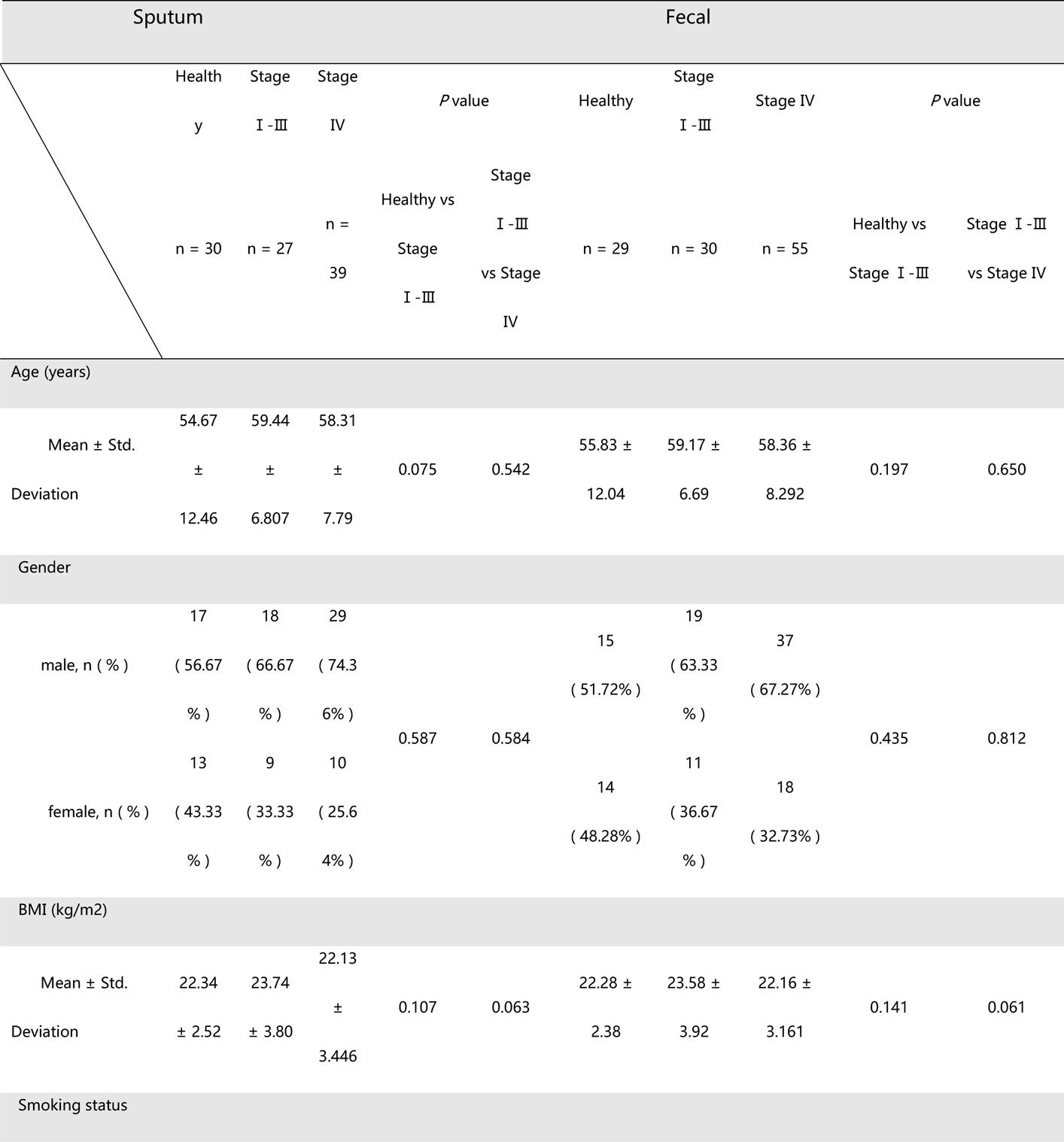
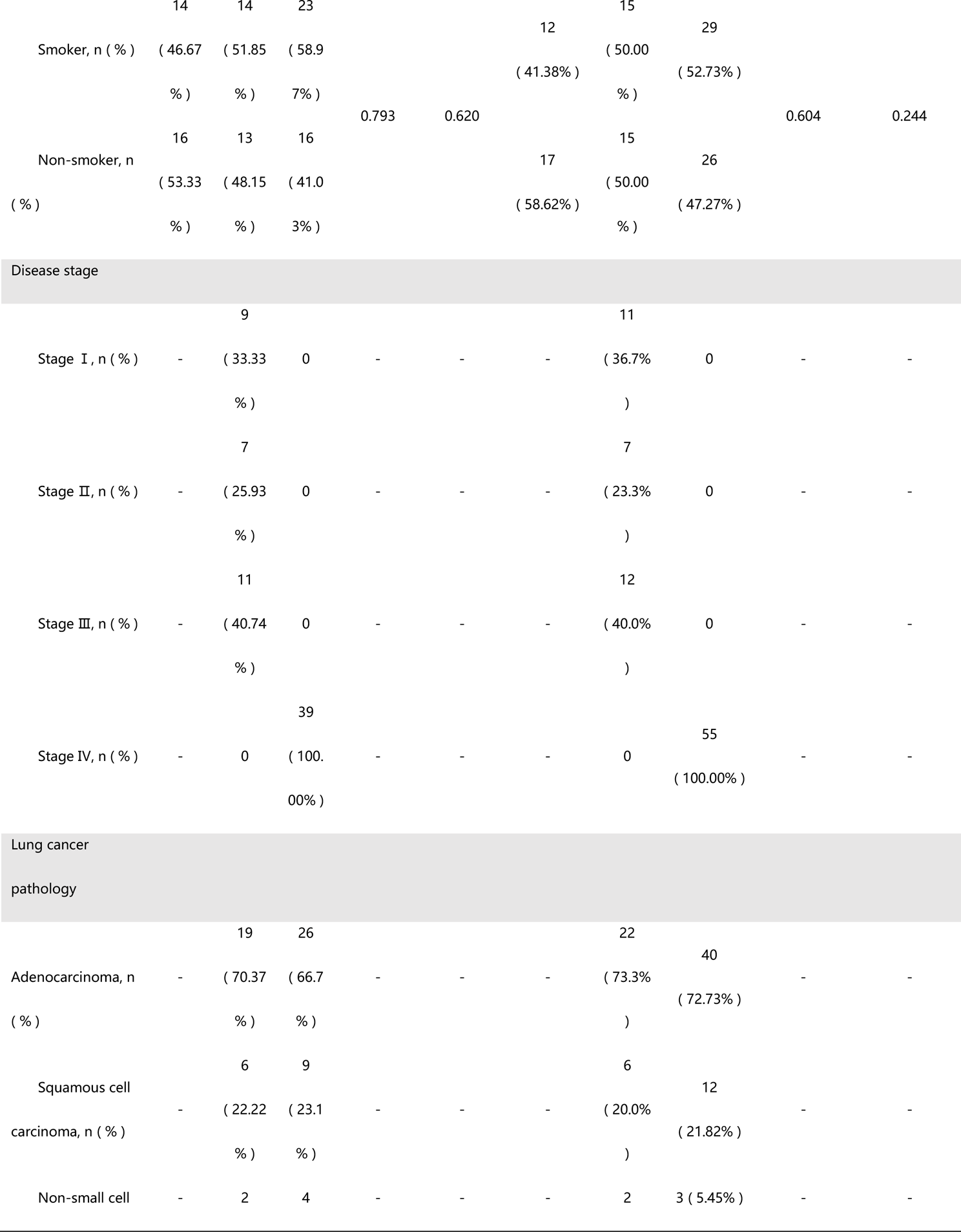
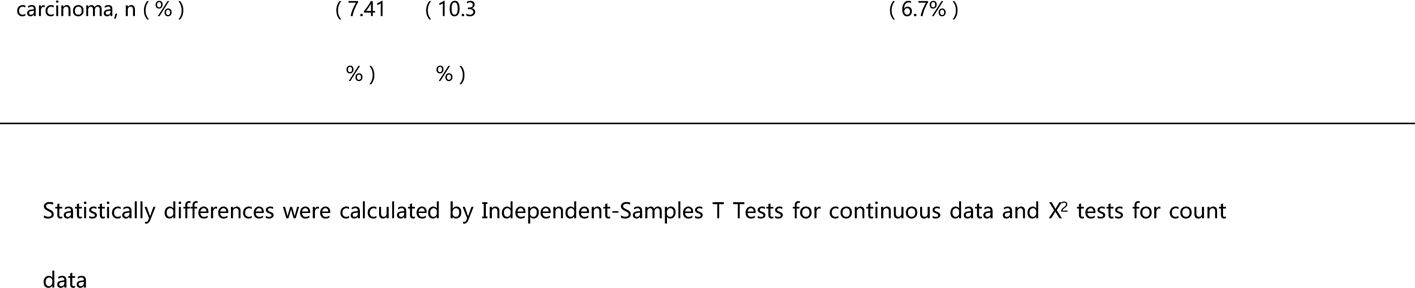
Clinical characteristics of healthy subjects and NSCLC patients.

We collected in total 30 sputum and 29 fecal samples from the healthy controls (Control) and 66 sputum and 85 fecal samples from the patients (NSCLC; see Figure 1A), and submitted them for 16S sequencing (see Methods). As shown in Figure 1 and Supplementary figure 1, we found that the microbial diversity, as measured by Shannon index, was significantly higher in sputum than in gut in the healthy controls as well as different disease stage groups (Figure 1B left panel; Supplementary figure 1A-B; Wilcoxon rank-sum test). We also performed principal coordinate analysis (PCoA) based on Bray-Curtis distance at genus level to assess the beta diversity in microbial composition and found that the sputum microbiota were significantly different from the gut in healthy controls (Figure 1B right panel) and patients of different disease stages (Supplementary figure 1A-B). Together, our results suggested that sputum microbiota were significantly different from the gut microbiota and had significantly higher microbial diversity.

**Figure 1.**
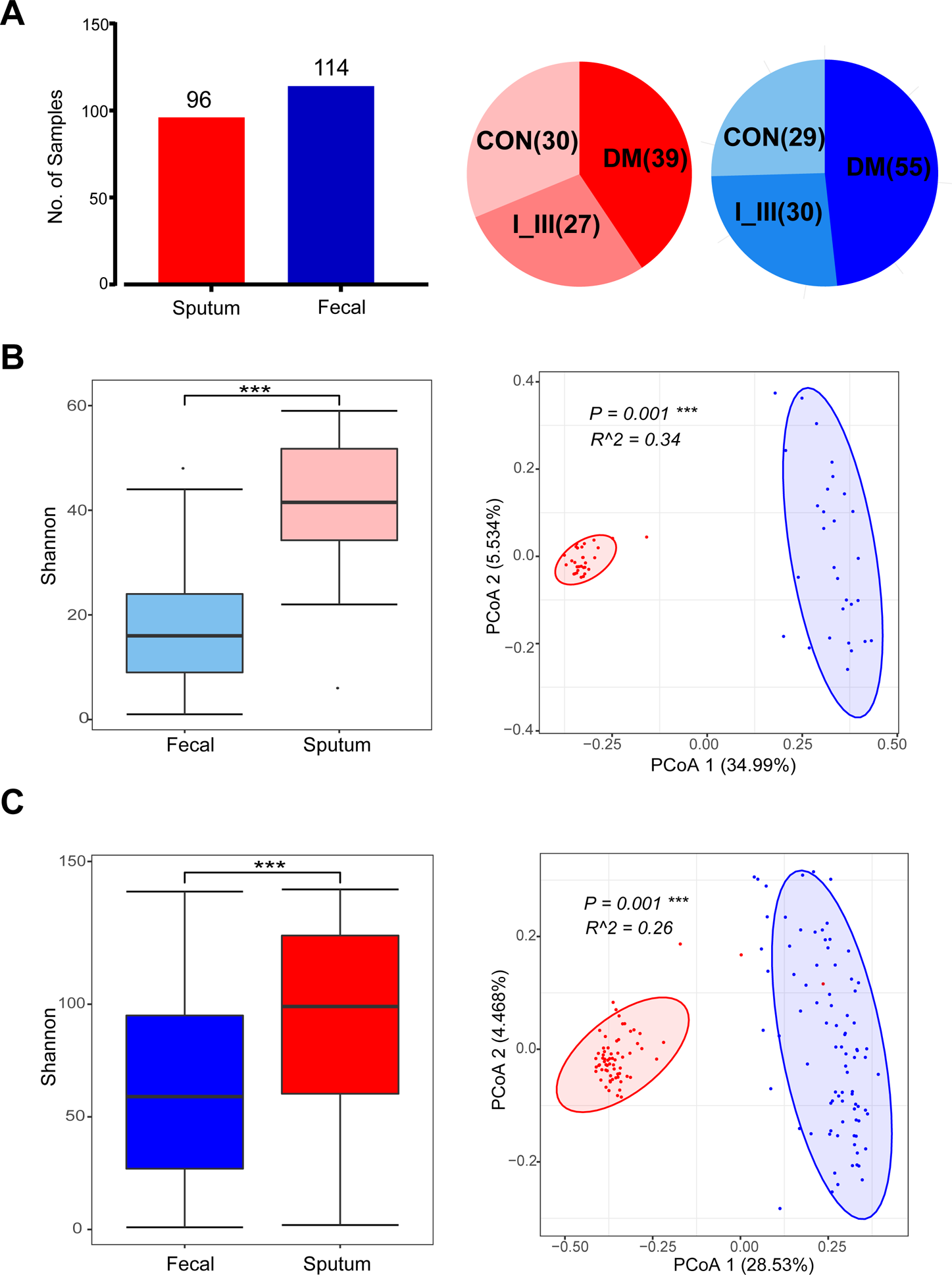
Sputum and gut microbiota differed significantly in terms of alpha- and beta-diversities. (**A**) Numbers of sputum (red) and gut (blue) samples collected in this study and their distributions in healthy controls and distinct disease stage groups. CON: healthy controls; I_III: patients with stages of I to III; DM: patients with distant metastasis (also referred to as stage IV). Disease stages were assigned according to the 8th American Joint Committee on Cancer (AJCC) guidelines [20]. (**B**) Comparisons of alpha-diversity and beta-diversity between sputum with gut in healthy controls. Shannon diversity index (alpha-diversity; left panel) was significantly lower in fecal; principal coordinate analysis (PCoA; right panel) based on Bray-Curtis distance at genus level showed that the overall microbiota composition was different between fecal and sputum samples. Wilcoxon rank sum tests were used to compare between groups. Level of significance: *** *P*<0.001; ** *P*<0.01; * *P*<0.05; NS. *P*≥0.05. (**C**) Comparisons of alpha-diversity (left panel) and beta-diversity (right panel) between sputum with gut in NSCLC patients (stages I to IV).

### Global alterations of sputum and fecal microbiotas in NSCLC patients of different stages

We next investigated the global alterations (i.e. dysbiosis) of sputum and gut microbiota in patients of different stages and between patients and healthy controls. As shown in Figure 2A, in the sputum microbiota, we found significant lower alpha-diversities (Shannon Index, left panel; Richness Index, middle panel) in NSCLC than the Control group. We also found that significantly different beta-diversities between NSCLC and Control (*P* = 0.001; Figure 2B, left panel) and between I_III and DM (*P* = 0.002; Figure 2B, right panel). Thus, the dysbiosis of sputum microbiota was associated with both NSCLC and the distant metastasis (stage IV).

**Figure 2.**
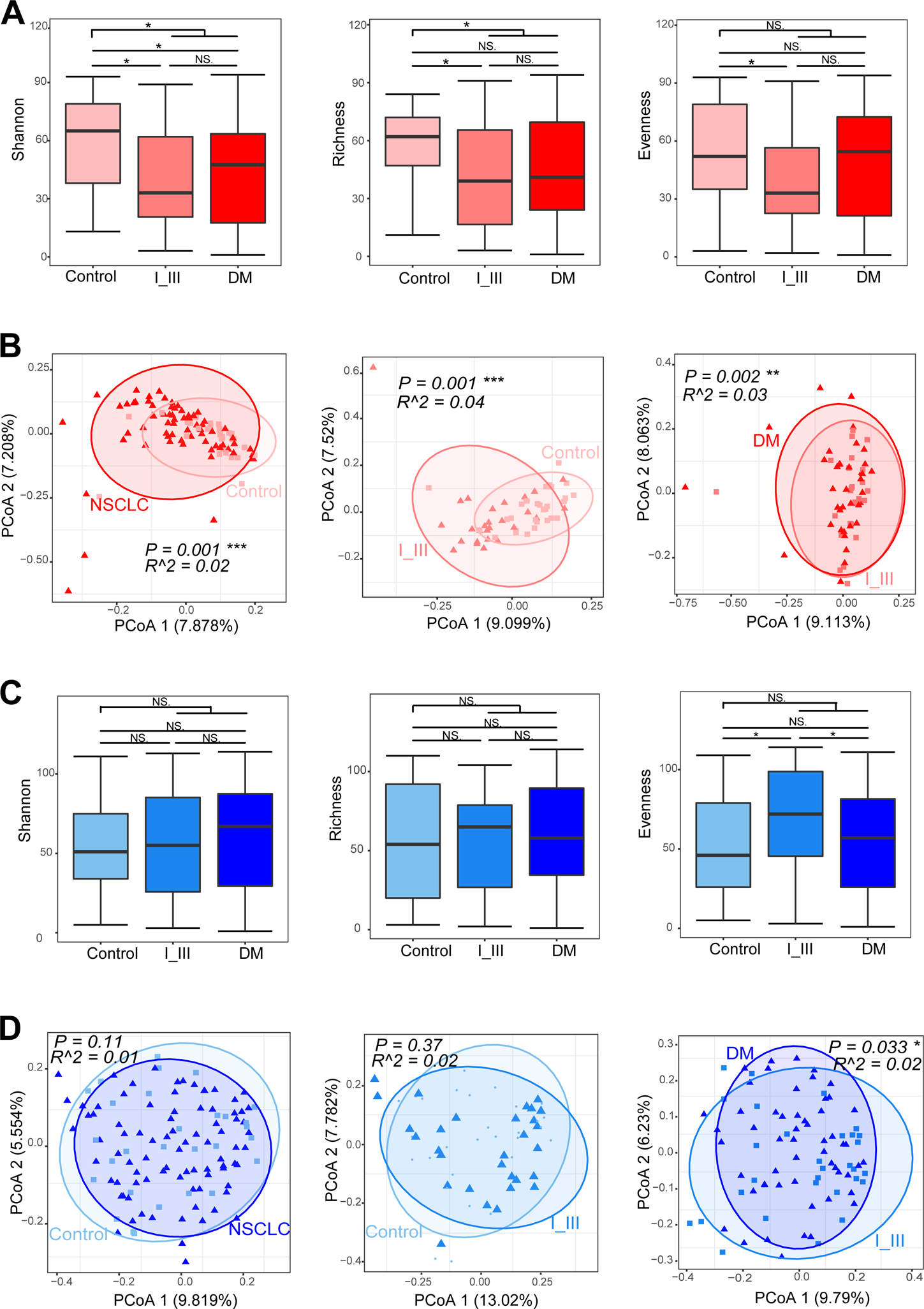
Global alteration of sputum microbiota was associated with NSCLC and distant metastasis (A-B), while fecal microbiota was only significantly associated with the latter (C-D). (**A**) Alpha diversity of sputum dysbiosis in pairwise comparisons. Shannon index (left); Evenness index (middle); Richness index (right). Shannon index and Richness index were significantly lower in patients as compared with controls; no significance was found in Evenness index. Wilcoxon rank sum test was used to compare between groups. Level of significance: *** *P*<0.001; ** *P*<0.01; * *P*<0.05; NS. *P*≥0.05. (**B**) Significant differences were found in beta-diversity between controls and NSCLC (left), as well as between controls vs I_III (middle) and I_III vs DM (right), indicating that dysbiosis of sputum microbiota was associated with lung cancer development and metastasis. Conversely, applying similar analyses to fecal samples, no alpha-diversities (**C**) but the beta-diversity in I_III compared with DM (**D**) was significantly different, suggesting that fecal microbiota dysbiosis was associated with distal metastasis, but not NSCLC.

Conversely, in the gut microbiota, we did not find significant differences between NSCLC and Control (Figure 2C) in neither alpha-diversities nor beta-diversities (Figure 2D, left panel). However, we found significant beta-diversities between I_III and DM patients (*P* = 0.033; Figure 2D, right panel); in addition, the microbial composition of DM was significantly different from I_III at genus level, with a decreasing evenness (Figure 2C, right panel). Together, the dysbiosis of the fecal microbiota was associated with distant metastasis, but not NSCLC.

### A significant proportion of microbial biomarkers was shared by sputum and gut microbiota

We then searched for individual taxa that showed differential abundances between subject groups (also known as microbial biomarkers) using LEfSe analysis (Linear discriminant analysis Effect Size; see Methods for details), and summarized the results in Figure 3.

**Figure 3.**
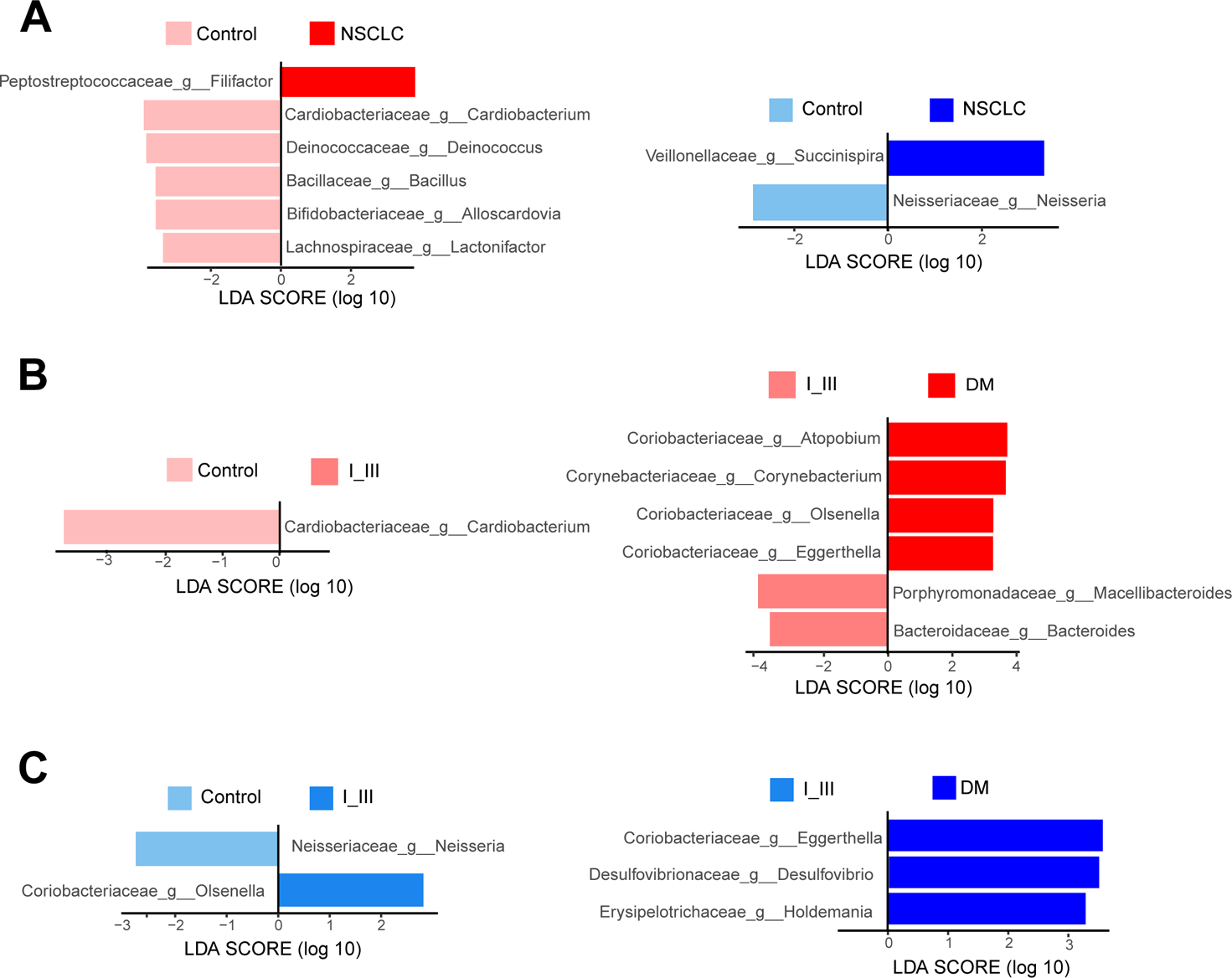
Shared and distinct microbial biomarkers between subject groups in sputum and feces microbiota. Differentially abundant microbial biomarkers between subject groups were identified using LEfSe analyses. (**A**) The relative abundance of 6 and 2 genera was significantly different between NSCLC and control group in sputum (left) and fecal (right), respectively. In order to identify biomarker for specific disease stages, we compared neighboring groups along the disease progression in sputum (**B**) and fecal (**C**). Control versus I_III, left; I_III versus DM, right.

We first compared all NSCLC patients as a whole (i.e. from stages I to IV) with the healthy controls. We found a genus, *Filifactor* was significantly enriched in NSCLC sputum samples (Figure 3A, left panel). *Filifactor* belongs to Firmicutes and contains a few pathogenic species (e.g. *F. alocis*) that are associated with periodontal diseases and endodontic lesions [21, 22]. This results suggested that *Filifactor* either represented part of the oral microbiota from the sampling, or could thrive as pathogens in other body sites like many other oral microbes did (e.g. *Fusobacterium nucleatum*) [23, 24]. Conversely, we found that a few genera, including *Cardiobacterium*, *Deinococcus*, *Bacillus*, *Alloscardovia* and *Lactonifactor* were depleted in sputum sample of the NSCLC group (Figure 3A, left panel). These results confirmed that the normal sputum microbiome has been significantly altered, since many of these genera were known members of healthy oral and/or gut microbiota [25, 26]. In addition, we found that the genus *Neisseria* was enriched in healthy controls and *Succinispira* was enriched in NSCLC patients in gut (Figure 3A, right panel). *Neisseria* belongs to the family *Neisseriaceae* and colonizes the mucosal surfaces of animals and contains a few known pathogenic species [27].

We next compared neighboring groups along the disease progression, i.e. Control versus I_III and I_III versus DM, in order to identify biomarker species for specific disease stages. We found that the *Cardiobacterium* was again identified to be enrichened in Control as compared with I_III (Figure 3B). In addition, we found a few biomarker species that were uniquely enriched in DM as compared with I_III, including three genera from the family *Coriobacteriaceae* (such as *Atopobium*, *Eggerthella*, and *Olsenella*). *Coriobacteriaceae* is a group of gram-positive bacteria that are often nonmotile, nonspore-forming, nonhemolytic and strictly anaerobic [28]. They are normal dwellers of mammalian body habitats including the oral cavity [29], the gastrointestinal tract [30], and the genital tract [31]. Consistent to our results, several members of the genera, including *Atopobium, Eggerthella, Gordonibacter, Olsenella,* and *Paraeggerthella* had been implicated in the development of various clinical pathologies including abscesses [32], periodontitis [33], intestinal diseases and tumors [34, 35]. Surprisingly, we found two genera of the family *Coriobacteriaceae* were identified as gut-biomarkers (Figure 3C). For example, genus *Olsenella* was also enriched in fecal samples of the I_III group as compared with the controls, while genus *Eggerthella* was also enriched the DM group as compared with I_III. Together, our results suggested that a significant proportion of sputum- and gut-microbial biomarkers were shared; the overlapping could be due to either extensive transmission from oral to other body sites [24], or the exposure to the same environment.

### The contributions of sputum and fecal microbiotas in patient stratification

We next assessed the potential value of sputum and gut microbiota in patient stratification. We generated predictive models using the Random forest algorithm implemented in Siamcat [36], evaluated the model performance with 10-times cross-validation and reported the averaged area under receiving operating characteristics curves values (AUROCs or AUC for short; see Methods) from 1000 repeats. We first generated models using the sputum and gut microbiota separately (referred to as sputum- and gut-models respectively). As shown in Figure 4A-D and Table 2, we found that sputum microbiota performed better than gut in patient stratification, in all subject group comparisons (Table 2).

**Figure 4.**
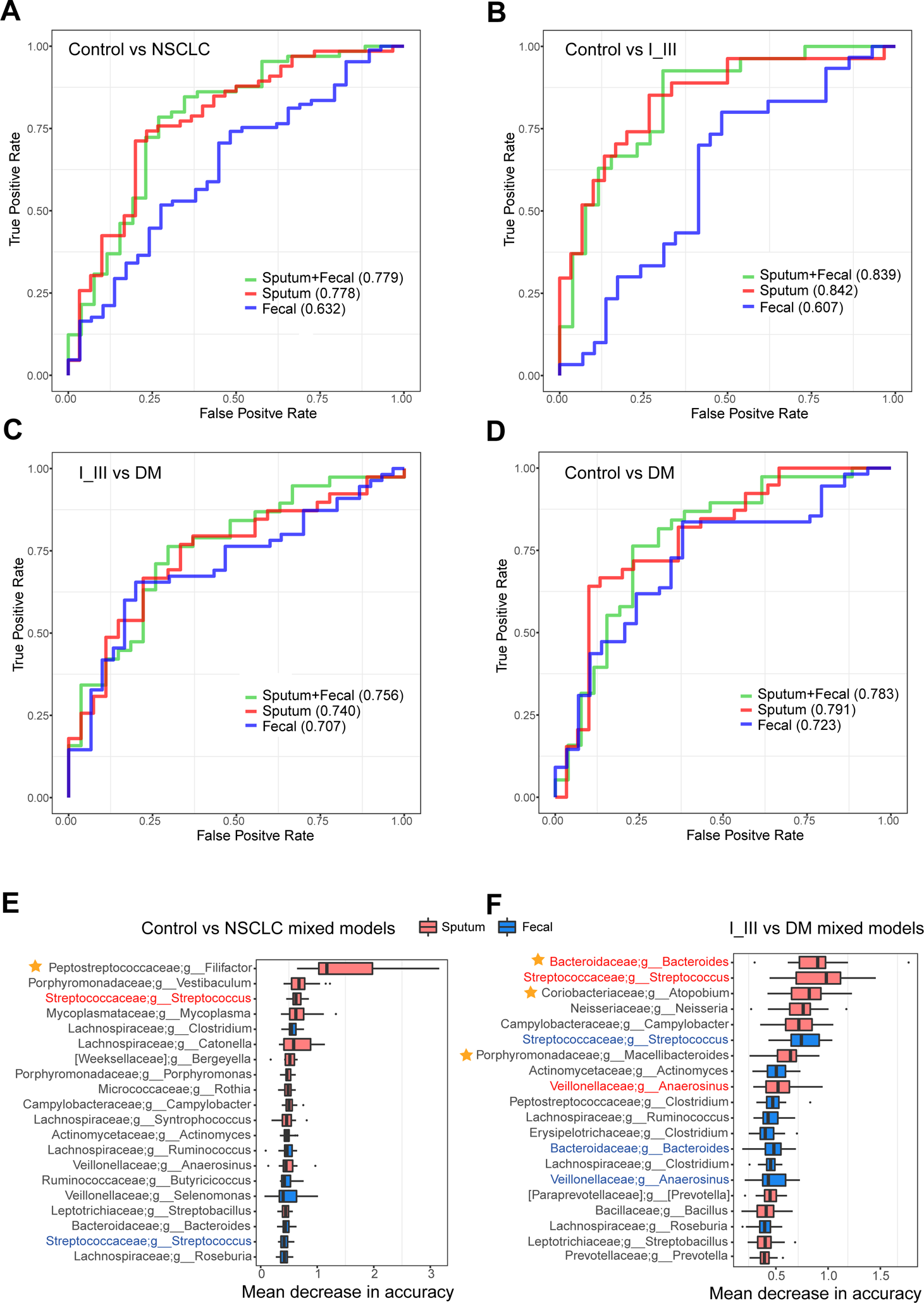
Disease classification based on taxonomic profiles of sputum, gut and both. Panels A to D showed the classification performance using relative abundance of genera as area under the ROC between subject groups. **(A)** Control vs NSCLC, **(B)** Control vs I_III, **(C)** I_III vs DM and **(D)** Control vs DM. Panels E to F showed the top twenty genera important to the mixed models; they were ranked by the median values of 1000 repeats, therefore boxplots were used here to demonstrate the medians and distributions of these values. **(E)** Control vs NSCLC and **(F)** I_III vs DM. Red boxes: sputum-derived genera; blue boxes: gut-derived genera. The colorful genera names indicated the overlap genera between sputum with gut. Star demonstrated the genus was significantly different in abundance using LEfSe analysis.

**Table 2.** The AUC values of classifying models.

|  | Control vs. NSCLC | Control vs. I_III | I_III vs. DM | Control vs. DM |
| --- | --- | --- | --- | --- |
| Sputum | 0.778 (0.673 - 0.883) | 0.842 (0.736 - 0.947) | 0.740 (0.618 - 0.862) | 0.791 (0.679 - 0.904) |
| Fecal | 0.632 (0.514 - 0.750) | 0.607 (0.458 - 0.756) | 0.707 (0.594 - 0.820) | 0.723 (0.608 - 0.838) |
| Sputum+Fecal | 0.779 (0.666 - 0.893) | 0.839 (0.730 - 0.948) | 0.756 (0.637 - 0.876) | 0.783 (0.661 - 0.905) |

We then built predictive models using both the sputum and fecal microbiome data as input (referred to as mixed models below). Among the enrolled subjects, we identified in total 91 subjects who had both sputum and fecal samples, among which 26, 27 and 38 were healthy controls, stage I_III and DM patients respectively. As shown in Figure 4A-D and Table 2, we found that the mixed model could perform either slightly better than or comparable to that of the sputum (Table 2).

We then examined the top twenty genera ranked according to their importance to the mixed models. As shown in Figure 4 & Supplementary Figure 2, there were more sputum-derived genera than gut-derived genera in numbers. For example, only seven and three gut-derived genera were among the top twenty in the Control versus NSCLC (Figure 4D) and Control versus I_III (Figure 4E) models, respectively. More importantly, the sputum-derived genera in general ranked higher in the mixed models and had higher cumulative importance scores (Table 3).

**Table 3.** The sum of cumulative importance scores in mixed models.

|  | Control vs. NSCLC | Control vs. I_III | I_III vs. DM | Control vs. DM |
| --- | --- | --- | --- | --- |
| Sputum | 8.030 (6.034 – 11.422) | 7.586 (5.235 – 10.690) | 7.230 (5.066 – 8.969) | 6.420 (5.416– 8.797) |
| Fecal | 3.277 (2.639 – 4.466) | 0.912 (0.618 – 1.366) | 4.599 (3.140 – 5.964) | 0.723 (0.608 - 0.838) |

Together, these results suggested that the sputum microbiota contributed more than the gut microbiota in patient stratification. In most cases, the sputum microbiota alone was sufficient for decent model performance.

### Top ranking taxa were also significantly shared by the sputum- and fecal-machine-learning models

We next checked if there were significant overlap in the top-ranking taxa between sputum- and fecal-models between controls and NSCLC; shared taxa often indicated that they may play similar roles at different body sites. As shown in Figure 5A-B, we found four of the top genera were shared at the same time in Control vs. NSCLC and Control vs. I_III models, including *Macellibacteroides*, *Streptococcus*, *Clostridium* and *Bacteroides*. *Bacteroides* maintained a complex and generally beneficial relationship with the host when retained in the gut, but when they escaped this environment they could cause significant pathology, including bacteremia and abscess formation in multiple body sites [37]. *Clostridium* were associated with a range of human diseases [38], and currently under investigation and testing as antitumor agents, because they germinated only in hypoxic tissues (i.e., tumor tissue), allowing precise targeting and direct killing of tumor cells [39]. Five out of twenty genera (*Anaerosinus*, *Clostridium*, *Bacteroides*, *Actinomyces* and *Streptococcus*) were shared by sputum and gut models of I_III vs. DM (Figure 5C). The human digestive tract was the main habitat for *Anaerosinus* [37]. There were several types of *Streptococcus*, two of which caused most of the strep infections in human: group A and group B [40]. These results indicated common features of sputum and gut dysbiosis during disease development and metastasis.

**Figure 5.**
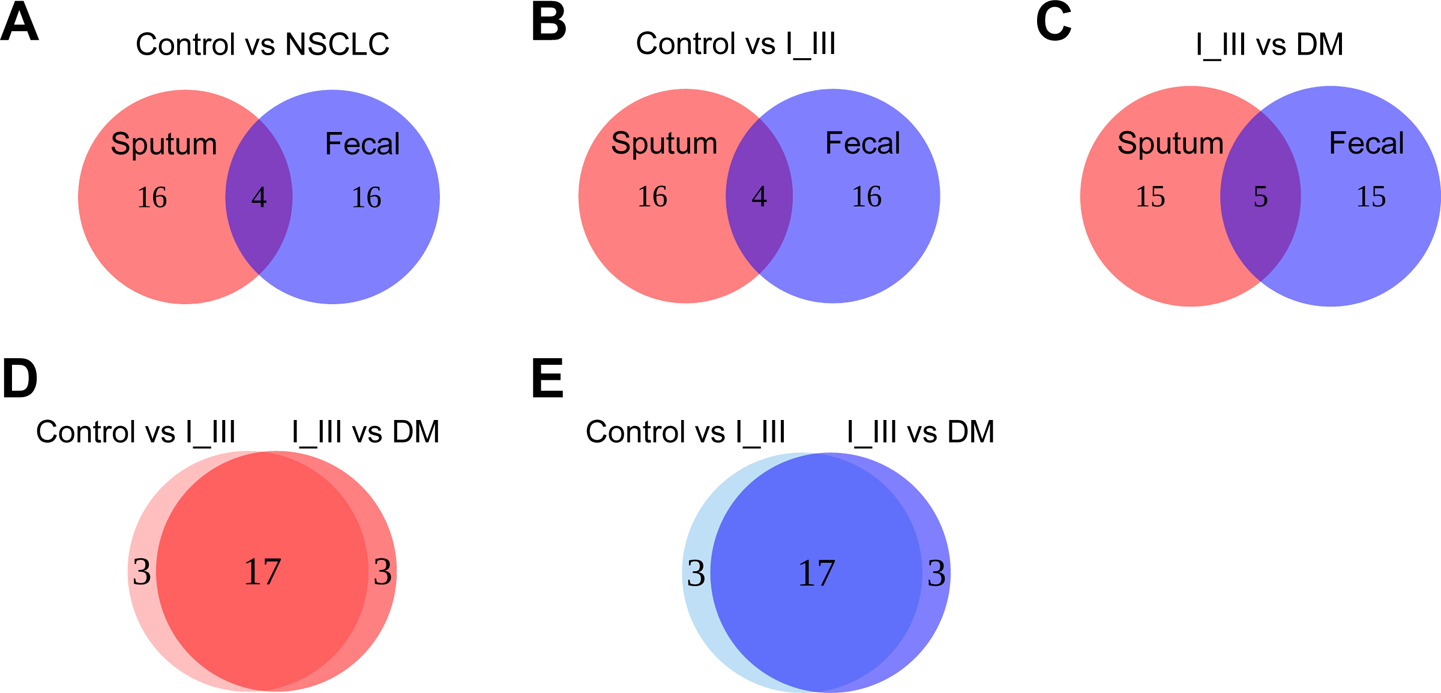
**Top-ranking genera in the machine learning models were significantly overlapped**. The Venn diagram showed the overlap of the top 20 genera between sputum with gut in (**A**) Control vs NSCLC classification model, (**B**) Control vs I_III classification model and (**C**) I_III vs DM classification model. The Venn diagram showed the overlap of top 20 genera (**D**) between sputum classification models and (**E**) between fecal classification models.

We also checked the overlapping of the top-ranking taxa in models between neighboring disease stages, such as models for Control vs. I_III and I_III vs. DM. Again, we found even more shared taxa. For example, we found seventeen out of the top twenty genera were shared in the two models generated using individual microbiota (Figure 5E-F). Unlike the sputum with more variety genera, there were two main families in gut, *Ruminococcaceae* and *Lachnospiraceae*; most members of which were found in human or animal digestive tract [41]. Previous studies have noted that both of them were depleted in patients with cirrhosis [42], enriched during alcohol abstinence and inversely correlated with intestinal permeability [43, 44]. These bacteria were known to have a beneficial effect on gut barrier function [44]. Not surprisingly, we found that in the mixed models, in which the same taxa from sputum- and fecal- were treated as distinct features, several of the above-mentioned taxa from both sputum and feces were among the top twenty taxa, including *Streptococcus* in the Control vs. NSCLC models, *Anaerosinus, Bacteroides* and *Streptococcus* in the I_III vs. DM models. Together, these results indicated that the same set of microbial taxa were underlying the development and progression of NSCLC, and the biomarkers for DM might be acquired early.

### *Pseudomonas aeruginosa*, a species implicated in infections, was enriched in brain-metastatic patients

Brain-metastasis (BM) represented the deadliest form of distant metastasis of NSCLC. To identify putative microbial biomarkers that were capable of distinguishing BM from other types of distant metastasis, we divided stage IV patients into two groups, namely the BM group (18 sputum samples and 25 fecal samples) and nonBM group (21 sputum samples and 30 fecal samples) (Figure 6A, left panel). As shown in Figure 6A, in the sputum microbiota, we found significantly different beta-diversities (*P*=0.011; middle panel) between the two groups, while there was no significant difference in fecal microbiota (*P*=0.178; right panel). Thus, the dysbiosis of sputum microbiota was in stronger association with brain metastasis of NSCLC than fecal. We next performed LEfSe analysis and Wilcoxon rank-sum test to identify potential microbial biomarkers between BM and nonBM groups (Figure 6B-C). Several differentially abundant genera were identified, including *Pseudomonas*, *Actinomyces* in sputum and *Blautia* and *Pseudomonas* in feces. *Pseudomonas* was highly abundant in the sputum of the BM group (∼8.14%) but not detectable in the nonBM group with relative abundance close to zero (Figure 6B, right panel); *Pseudomonas* was also not detectable in any other disease stages nor in healthy controls. *Pseudomonas* was also significantly enriched in fecal samples of the BM group (with relative abundance of ∼0.47%) and not detectable in other fecal samples.

**Figure 6.**
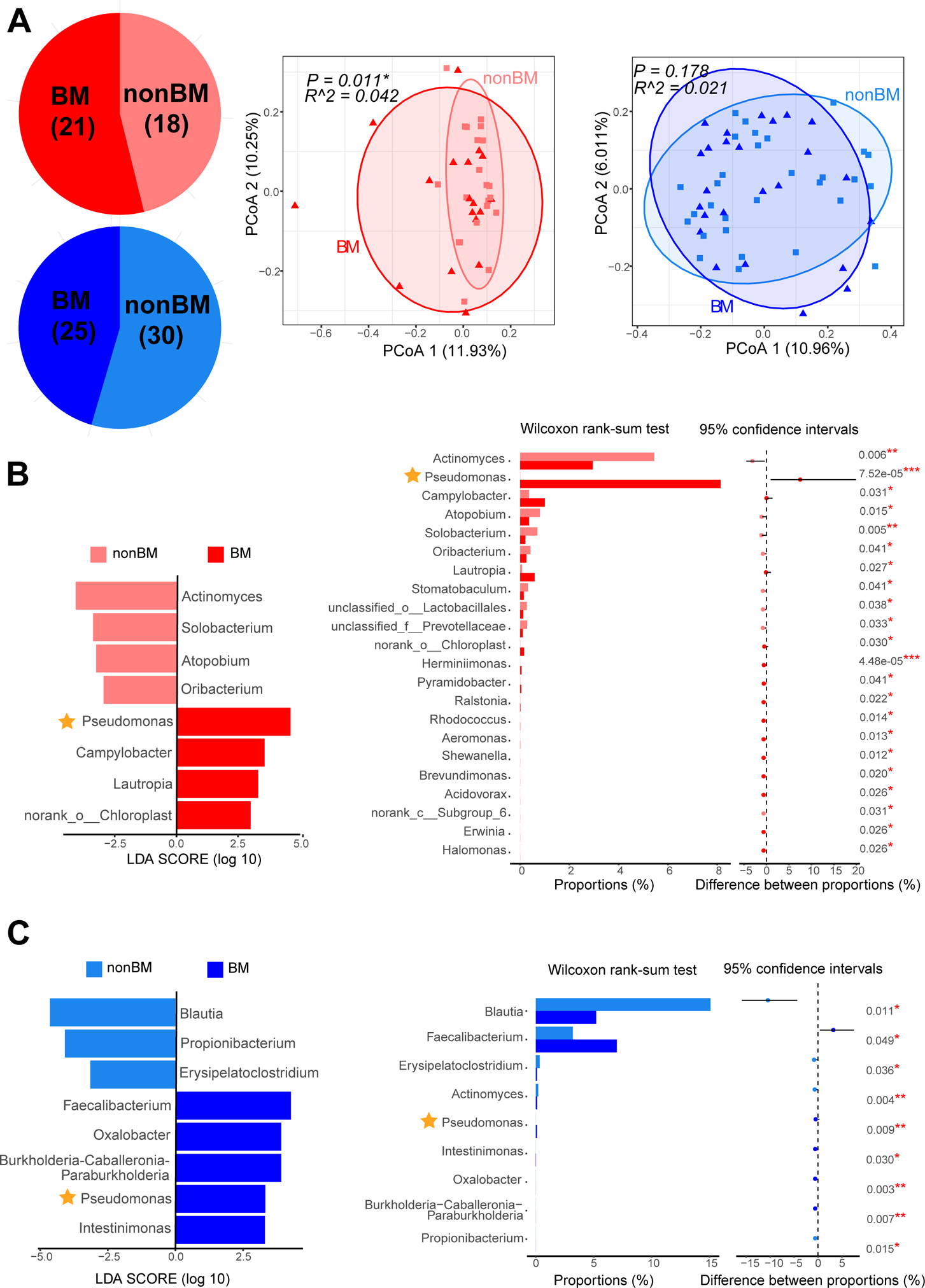
Patients with brain metastasis differed significantly from other distant metastasis in microbial profiles of sputum and feces. (**A**) Numbers of sputum (red) and gut (blue) brain metastasis samples (left). BM: NSCLC patients in stage IV with brain metastasis, nonBM: stage IV NSCLC patients without brain metastasis. PCoA analysis showed differences on beta-diversity between BM and nonBM in sputum (middle) not gut (right). LEfSe (left) analysis and Wilcoxon rank-sum test (right) of differentially abundant microbial biomarkers between BM and nonBM in sputum (**B**) and gut (**C**). Level of significance: *** *P*<0.001; ** *P*<0.01; * *P*<0.05; NS. *P*≥0.05. Star demonstrated the genus *Pseudomonas* was significantly different in abundance.

We then generated the distinguishing BM and nonBM models using the sputum microbiota, gut microbiota and mixed microbiota separately. As shown in Figure 7A, we found that sputum microbiota performed best in BM and nonBM group comparison. We also examined the top-ranking taxa in sputum-, fecal- and mixed models. As shown in Supplementary Figure 3, there were more sputum-derived genera than gut-derived genera in numbers. Only three gut-derived genera were among the top twenty in the BM versus nonBM mixed model. Again, we found *Pseudomonas* was the most important genus to sputum- and mixed models between BM and nonBM (Figure 7B and Supplementary figure 3). Thus, *Pseudomonas* is a prominent biomarker for brain metastasis in sputum. *Pseudomonas* consists of a groups of aerobic, Gram-negative and rod-shaped bacteria [1] that are associated with many human diseases but are relatively rare in the healthy gut (see https://gmrepo.humangut.info/species/286 for an overview their prevalence and abundances in gut microbiota associated with human health and diseases [45]). According to a MAPseq tool [46], which assigns 16S sequencing reads to distinct taxa with confidence scores, most of the *Pseudomonas* reads could be reliably identified as *Pseudomonas aeruginosa* (see Methods for details)*. P. aeruginosa* is one of the major causes of nosocomial infections worldwide [3] and is often associated with long-term wounds, pneumonia [4], chronic obstructive lung diseases [47], cystic fibrosis explanted lung [5], bronchiectasis [48] and chronic destroyed lung disease due to tuberculosis [47]. Its roles in brain metastasis needs to be further explored.

## Discussion

We believed that the present study is the first to investigate the alterations of both sputum (as a proxy for lung) and gut microbiota on the development and metastasis of NSCLC. The results of our study suggest that lung microbiota may play major roles in the development of NSCLC, the dysbiosis of which could accurately stratify patients from healthy controls, while the distant metastasis (DM) was associated with both sputum and gut microbiota dysbiosis. We further identified a prominent microbial biomarker for brain metastasis (BM).

In recent years, growing evidence have linked the alterations in lung or gut microbiota to LC or NSCLC. However, the relative importance of the gut and lung microbiota to the development of NSCLC are still unclear; in addition, their alterations along with DM of NSCLC have not been characterized. Therefore, in this study we assembled a cohort including patients of diagnosed NSCLC, including those suffered from DM (stage IV), and collected both sputum and fecal samples. We delineated the microbial community structure by 16S rRNA sequencing. The sputum and gut microbiota differed significantly in terms of alpha-diversity and beta-diversity, regardless health statuses and disease stages; surprisingly, sputum microbiota had significantly higher richness (taxon count) and evenness than gut microbiota, suggesting unappreciated microbial complexity in the respiratory systems and putative important roles in related diseases. We built machine learning models to evaluate the relative importance of sputum and gut microbiota in patient stratification. We found that both sputum and gut microbiota dysbiosis contributed significantly to discriminating metastatic to non-metastatic patients, while sputum microbiota performed the best in discriminating stage I_III patients from healthy controls. These results highlighted the potentials using both sputum and gut microbiota in non-invasive disease diagnosis.

By comparing to healthy controls of matching demographic and clinical characteristics, we identified microbial biomarkers that showed significant abundance differences between subject groups. Not surprisingly, many of the identified biomarkers were either previously associated with other diseases [38, 40], or known to induce inflammation and/or interact with host immunity [31–38]. For example, the genera *Atopobium, Eggerthella* and *Olsenell* (Figure 3C,F), belong to the family *Coriobacteriaceae*, had been implicated in the development of various clinical pathologies including abscesses [32], periodontitis [33], intestinal diseases and tumors [34, 35]; and that *Atopobium* was the third important genus to I_III vs IV mixed model (Figure 4F). Similarly, a genus *Filifactor*, which was the most important genus in the Control vs NSCLC mixed model, was significantly enriched in NSCLC patients; it was known that some species of *Filifactor* were members of human oral microbiome and were pathogenic [21].

We found significant overlap between sputum- and fecal-biomarkers, suggesting that these microbes may play similar roles at different body sites. In addition, most of the microbial-biomarkers of distinct disease stages, i.e. I_III vs. healthy controls and DM vs. I_III, also overlapped (Figure 5); we found that the cumulative abundances of these biomarkers were increased (decreased) continuously along disease development. These results suggested that distant metastasis (DM) was the ultimatum development of lung cancer, and the DM-modulating microbes were acquired early.

We identified *Pseudomonas aeruginosa* as a prominent biomarker for brain metastasis (BM); *P. aeruginosa* was highly abundant in BM patients as compared with other NSCLC as well as other distant metastatic patients and was exclusively found in sputum. *P. aeruginosa* is found in many diseases and is often associated with long-term wounds; its role in BM should be further experimentally determined.

Despite the strengths of our study, there were two limitations. First, currently only limited numbers of subjects were enrolled, which could limit the predictive performance of our patient stratification models; better ML models would have been possible with more subjects and deeper coverage of metagenomics sequencing data. Second, the exact roles of gut and lung microbiota in NSCLC and metastasis needed to be further illustrated. Further experiments are needed to investigate their relative contributions by removing one at a time.

### Conclusions

In summary, we surveyed both sputum (as a proxy for lung) and gut microbiota of patients with NSCLC and distant metastasis and compared them with healthy controls. We obtained mathematical models capable of distinguishing patients from healthy controls as well as patients at different disease stages with high performance. The top taxa ranked by these models could be used for future experiments to illustrate the underlying molecular mechanisms, and/or biomarkers for disease diagnosis. Our analyses revealed that the alterations of sputum (as a proxy to lung) microbiota have stronger association with NSCLC and distant metastasis than the gut, indicating that tumor-site associated microbiota may contribute more to disease development.

## Methods

### Study design and sample collection

NSCLC patients were recruited in the Cancer Center, Union Hospital, Tongji Medical College, Huazhong University of Science and Technology, China. Healthy relatives of these patients were recruited as healthy subjects. The criteria for selecting controls were as following: good physical status, no significant respiratory or alimentary conditions. NSCLC diagnosis was established according to histological criteria. Clinical stage of NSCLC was determined following the 8th American Joint Committee on Cancer (AJCC) guidelines [20]; patients were classified into four distinct disease stages (i.e. from I to IV), in which stage IV referred to distant metastasis. No distant metastasis to any regions of the intestines was collected in this study.

The main exclusion criteria were as following: less than 18 years of age; any antibiotic therapy within the previous 1 month; known COPD (chronic obstructive pulmonary disease), pneumoconiosis, silicosis or any other diseases of the respiratory system; inability to give written informed consent. This study was approved by the Ethical Committees of the Cancer Center and registered with ClinicalTrials.gov (Identifier: NCT 03454685). All participants provided written informed consent before sample donation.

All fecal and spontaneous sputum samples were obtained after NSCLC diagnosis and before the patients received treatment. These samples were immediately placed in −80 ℃. Demographic and clinical data, including smoking status, gender, age, body mass index (BMI), disease stage and lung cancer pathology were obtained from each participant.

### DNA Extraction

Bacterial DNA was extracted from the fecal and sputum samples using the OMEGA-soil DNA Kit (Omega Bio-Tek, USA) according to the manufacturer’s instructions. The quality of DNA was measured using a NanoDrop 2000 Spectrophotometer (Thermo Scientific, USA). The quality of DNA was detected by 1% agarose gel electrophoresis. Bacterial DNA was immediately stored at −80 °C until further analysis.

### 16S rRNA amplification and sequencing

Bacterial DNA was isolated from fecal and sputum samples as previous described. DNA libraries covering the V3-V4 hypervariable regions of the bacterial 16S-rDNA gene were constructed using the FastPfu Polymerase (TransGen, China) according to the manufacturer’s instructions. We used the primer set composed of 338F: 5’ – ACTCCTACGGGAGGCAGCAG-3’, and 806R: 5’ – GGACTACHVGGGTWTCTAAT-3’, which was designed to amplify the V3–V4 hypervariable region. All PCR products were purified with an AxyPrep DNA Gel Extraction Kit (Axygen Biosciences, USA) and quantified using a QuantiFluor™-ST (Promega, USA) according to the manufacturer’s instructions. The sequencing of the PCR amplification products was performed on an Illumina Miseq platform (Illumina, USA) by Majorbio Bio-Pharm Technology Co., Ltd. (Shanghai, China). Sequence data has been deposited to the NCBI SRA database under the NCBI bioproject ID PRJNA576323.

### Sequencing data analysis and taxonomic assignment

Overall read quality was checked for each sample using FastQC. After Trimmomatic, reads with quality less than 30 or length less than 100 bp were removed from subsequent analysis. The filtered reads were then analyzed using Qiime2 (version 2018.11) [49]. DADA2 software, wrapped in QIIME2, was used to filter the sequencing reads and construct feature table. The taxonomy classify database was downloaded from Qiime2 (gg-13-8-99-515-806-nb-classifier.qza). Taxa with relative abundance less than 0.001 was removed. All analyses were carried out on genus level except for the alpha diversity. The taxonomy classify on species level was identified using “MAPseq” [46], which is a highly efficient approach with confidence estimates, for reference-based rRNA analysis; while also providing more accurate taxonomy classifications.

### Statistics analysis

Patients’ characteristics were expressed as mean ± std. deviation and compared using Χ2 tests or Independent-Samples T Test as appropriate. Statistical analyses were performed using SPSS V.19.0 for Windows (Statistical Product and Service Solutions, Chicago, Illinois, USA).

The beta diversity analyses were performed using the R package “Vegan”. Principal coordinate analysis (PCoA) and adonis analysis were performed based on Bray-Curtis distance. Linear discriminant analysis effect size (LEfSe) analysis [50] and Wilcoxon rank-sum test [51] were used to identify differentially abundant genera between subject groups. R package “Siamcat” [36] was used for Random forest modeling and 10-fold cross validation with 100 times repeat. The operating characteristic curves (receiving operational curve, ROC) were constructed and area under curve (AUC) was calculated to assess the diagnostic performance of the model with the pROC package [52].

### Availability of data and materials

Sequencing data is available and has been deposited to the NCBI SRA project under the NCBI BioProject ID PRJNA576323. Methods, including statements of data availability and additional references, are available at the publisher’s website.

## Acknowledgements

We are grateful for all the subjects who participated in the study.

## Funding

This work was partly supported by National Natural Science Foundation of China (81573090, 81773233, 61932008, 61772368, 61572363), National Key R&D Program of China (2018YFC0910500), Natural Science Foundation of Shanghai (17ZR1445600), Shanghai Municipal Science and Technology Major Project (2018SHZDZX01) and ZJLab.

### Author information

Hui Lu, Na L. Gao and Chunhua Wei contributed equally to this work.

### Affiliations

Cancer Center, Union Hospital, Tongji Medical College, Huazhong University of Science and Technology, 430074 Wuhan, Hubei, China

Hui Lu, Chunhua Wei, Fan Tong, Jiaojiao Wang, Huanhuan Li, Ruiguang Zhang, Hong Ma, Ye Wang, Zhiwen Liang, Hao Zeng & Xiaorong Dong

Key Laboratory of Molecular Biophysics of the Ministry of Education, Hubei Key Laboratory of Bioinformatics and Molecular-imaging, Department of Bioinformatics and Systems Biology, College of Life Science and Technology, Huazhong University of Science and Technology, 430074 Wuhan, Hubei, China

Na L. Gao & Wei-Hua Chen

Department of medical oncology, lung cancer and gastrointestinal unit, Hunan cancer hospital/The Affiliated Cancer Hospital of Xiangya School of Medicine, Central South University, Changsha, China, 410013

Nong Yang & Yongchang Zhang

Huazhong University of Science and Technology Ezhou Industrial Technology Research Institute, 436044 Ezhou, Hubei, China

Wei-Hua Chen

College of Life Science, HeNan Normal University, 453007 Xinxiang, Henan, China

Wei-Hua Chen

### Contributions

W.H.C. and X.R.D. designed the study. X.R.D., W.H.C., H.L., N.L.G. and C.H.W. designed the experiments. H.H.L., R.G.Z., H.M., N.Y., Y.C.Z., Y.W., and Z.W.L collected samples. H.L., C,H.W., J.J.W and F.T. performed the 16S-seq and clinical data. N.L.G. and H.Z. analyzed the sequencing data and performed statistical analyses. X.R.D, W.H.C., N.L.G. H.L. and C.H.W. wrote the manuscript with all authors contributing to the writing and providing feedbacks. All authors read and approved the final version of the manuscript.

### Corresponding authors

Correspondence should be addressed to Wei-Hua Chen or Xiaorong Dong.

### Ethics declarations

#### Ethics approval and consent to participate

This study was approved by the Ethical Committees of the Cancer Center and registered with ClinicalTrials.gov (Identifier: NCT 03454685); Cancer Center, Union Hospital, Tongji Medical College, Huazhong University of Science and Technology (2018-S271).

### Consent for publication

Not applicable.

## Additional information

Correspondence and requests for materials should be addressed to Wei-Hua Chen and Xiaorong Dong.

---

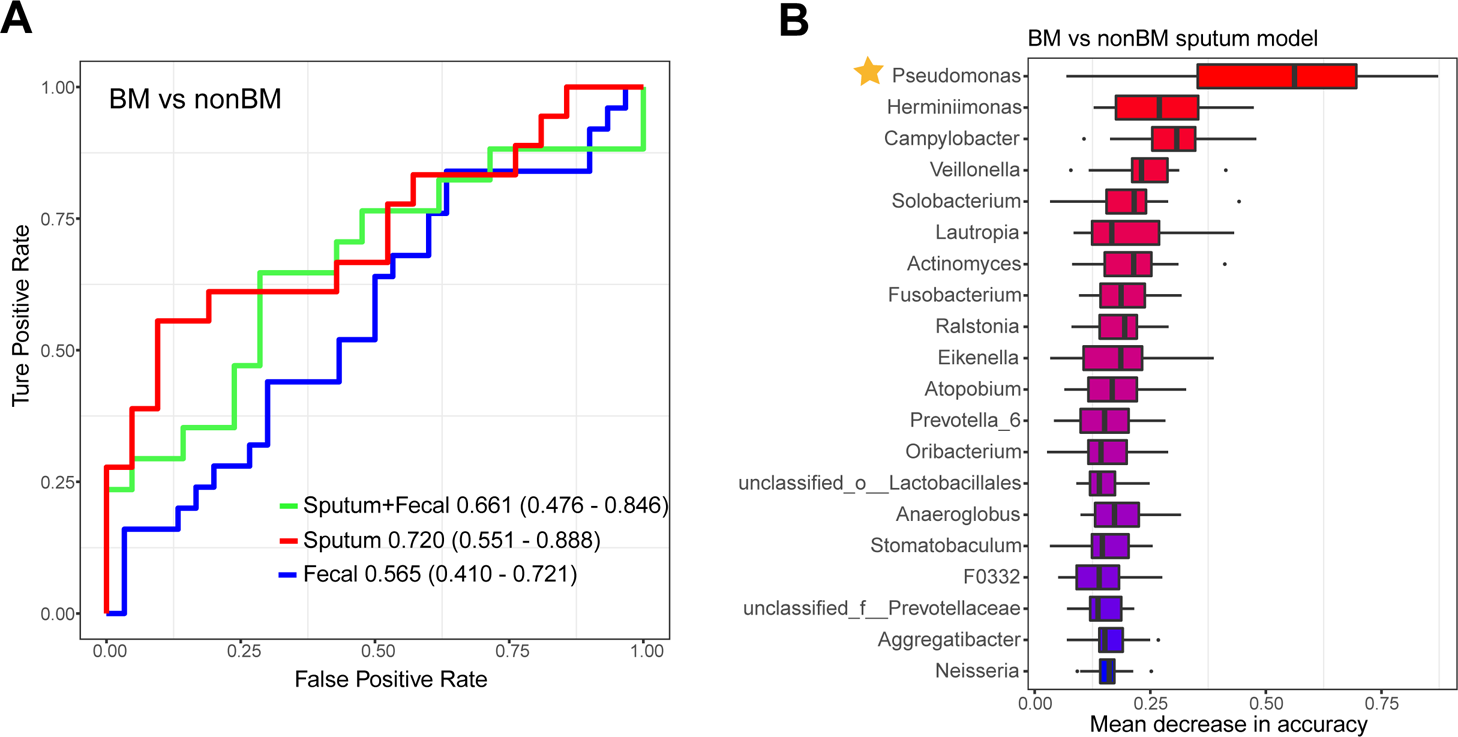

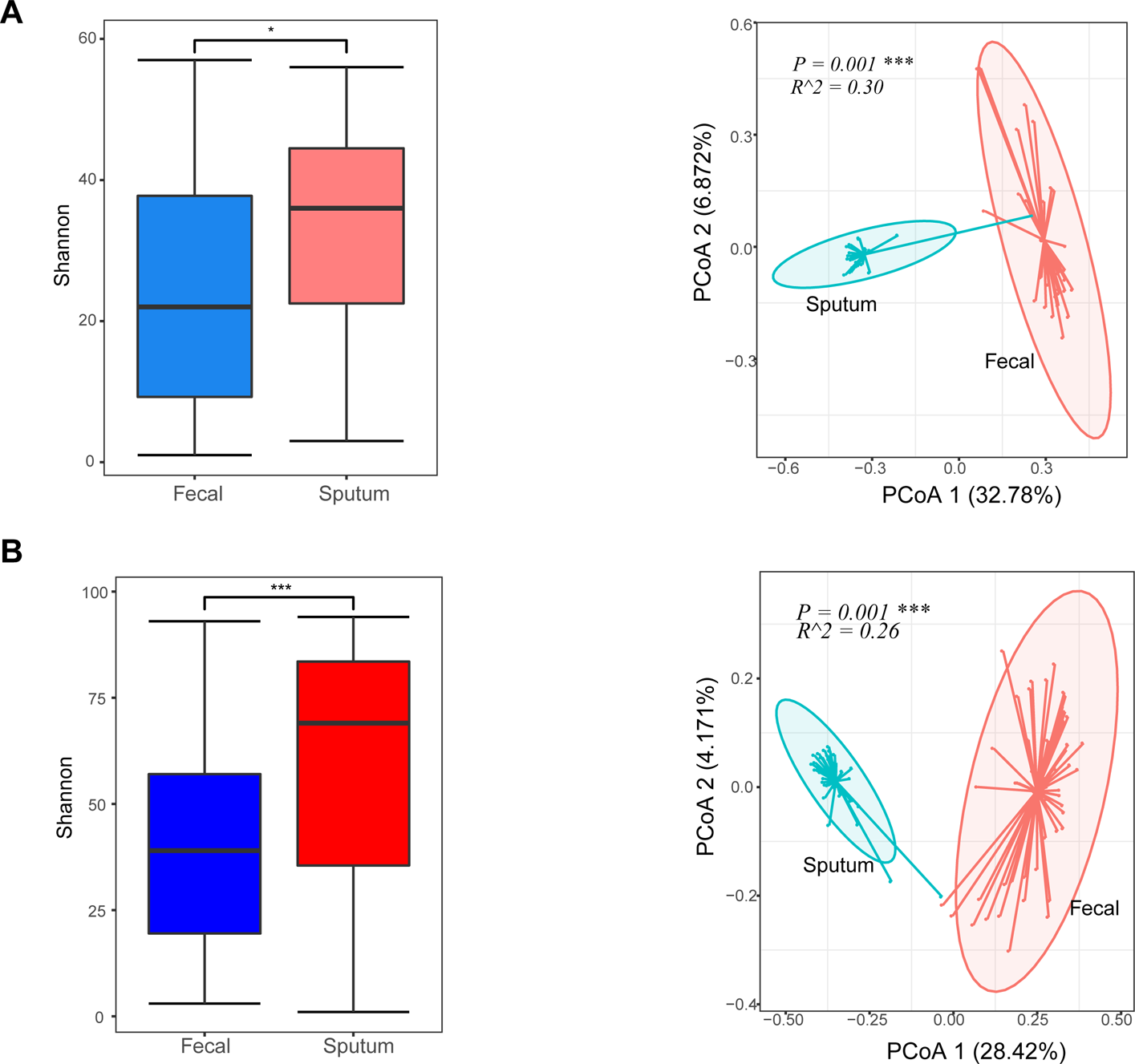

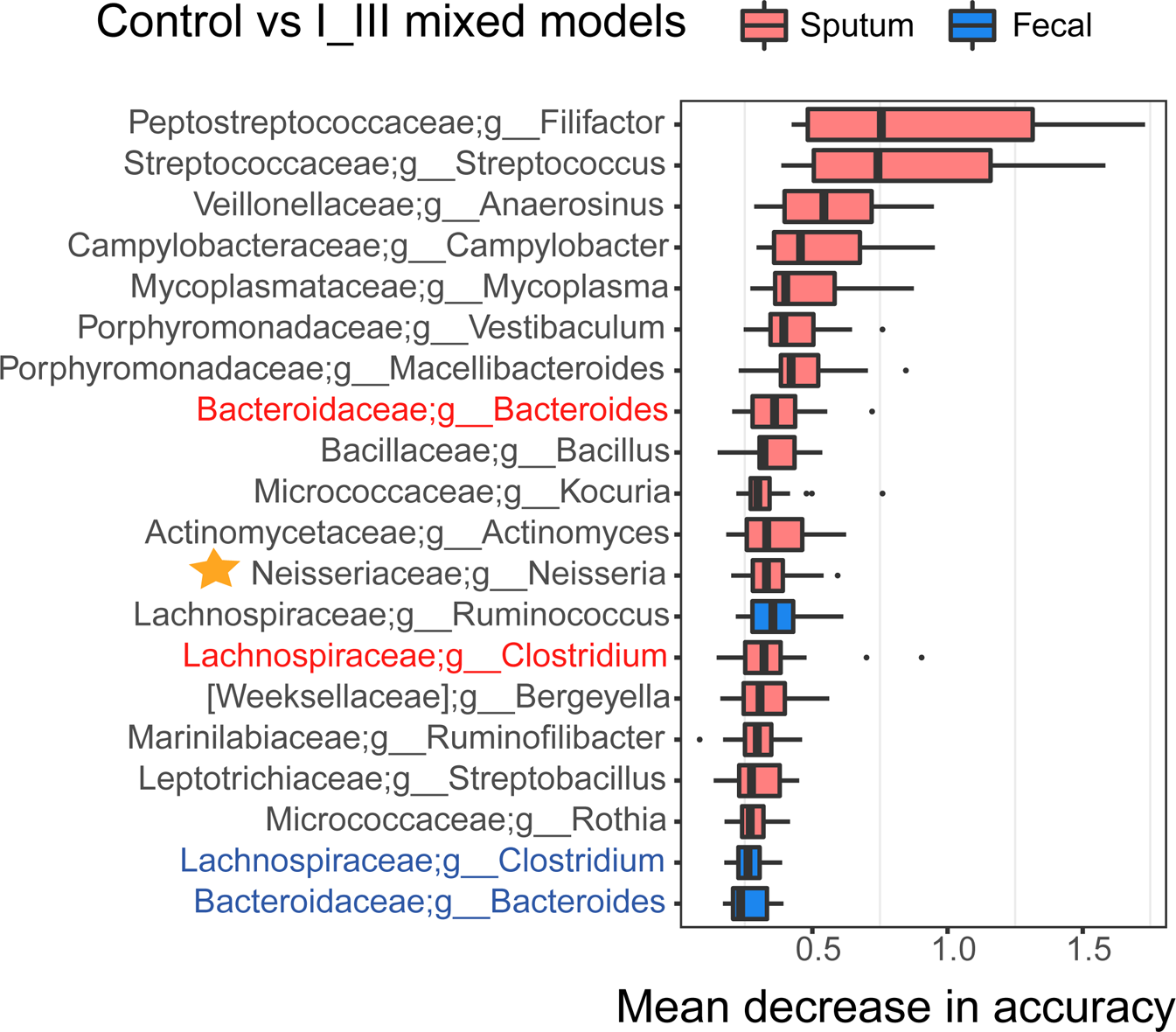

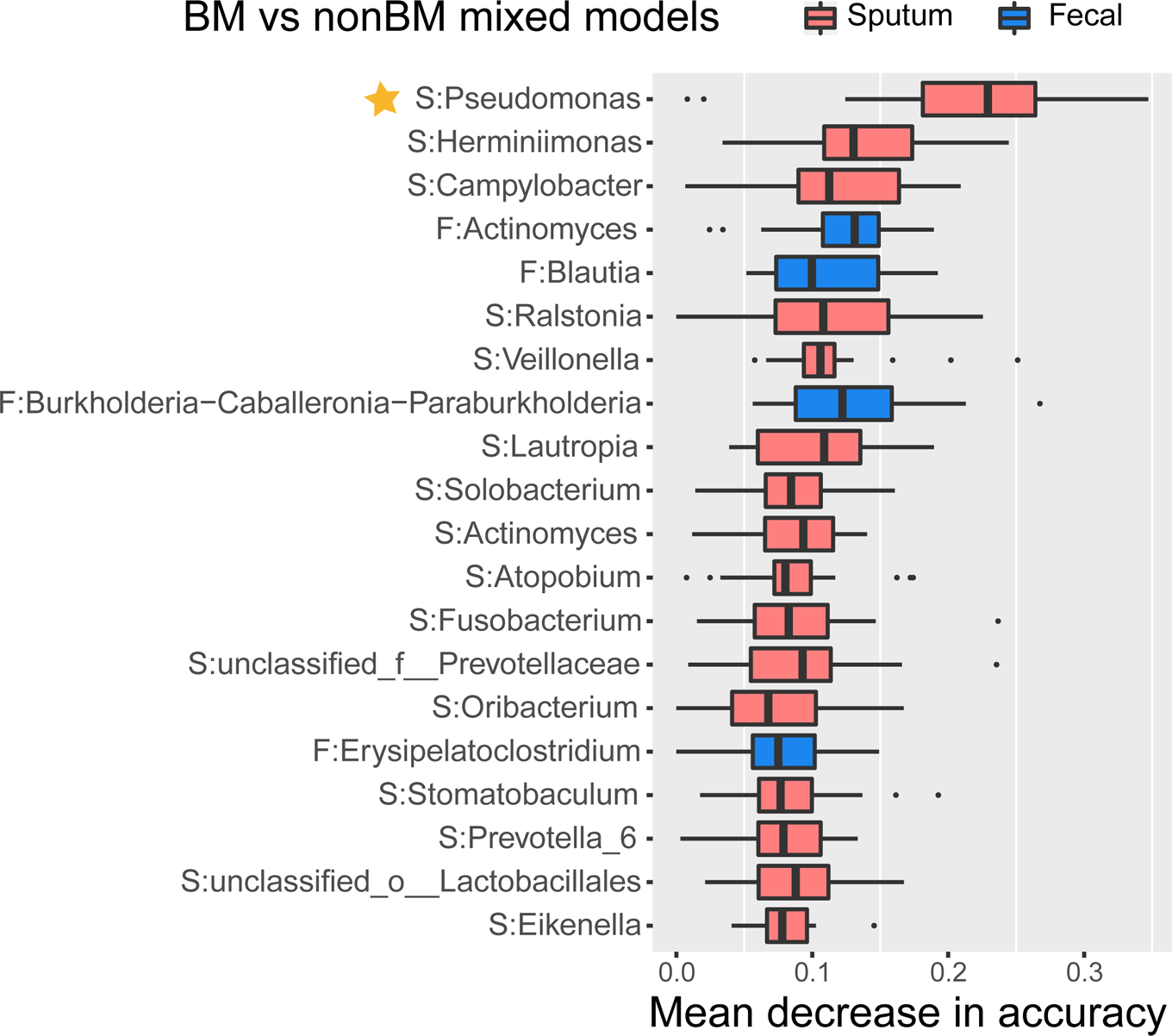

