## Supplementary1 for "Alterations of human lung and gut microbiome in non-small cell lung carcinomas and distant metastasis"

### Supplementary data

**Supplementary figure 1. Sputum and gut microbiota differed significantly in terms of alpha- and beta-diversities in patients of different disease stages.** (**A**) Shannon diversity index (alpha diversity) was significantly lower in fecal than in gut samples of I_III patients (left); PCoA showed that the overall microbiota composition was different between fecal and sputum in I_III patients (right). (**B**) Shannon diversity index was significantly lower in fecal than in gut samples of DM patients (left); PCoA showed that the overall microbiota composition was different between fecal and sputum in stage IV patients (right).

**Supplementary figure 2**. The top twenty genera ranked according to their importance to the mixed models in Normal vs I_III. Red boxes: sputum-derived genera; blue boxes: gut-derived genera. The colorful genera names indicated the overlap genera between sputum with gut. Star demonstrated the genus was significantly different in abundance using LEfSe analysis.

**Supplementary figure 3**. The top twenty genera ranked according to their importance to the mixed models in BM vs nonBM. Red boxes: sputum-derived genera; blue boxes: gut-derived genera. Star demonstrated the genus *Pseudomonas* was significantly different in abundance.
